# Distinct prostate cancer neuroendocrine subtypes predict prognosis and guide personalized prostate-specific membrane antigen-targeted therapy

**DOI:** 10.1101/2024.09.28.615561

**Authors:** Yung-Chih Hong, Cheng-Han Tsai, Tze-Yun Hu, Chih-Sin Hsu, Yu-Ching Peng, Weber Chen, William J. Huang, Tzu-Ping Lin, Pei-Ching Chang

## Abstract

**Background and Objective:** Second-generation hormonal therapy inhibits castration-resistant prostate cancer (CRPC), but the tumor eventually recurs as neuroendocrine prostate cancer (NEPC) and turns lethal. Differentiating lineage plasticity that contributed to distinct NEPC subtypes aids in advancing treatments, particularly the recent FDA-approved ^177^Lu-PSMA-617 radiopharmaceutical therapy.

**Methods:** We integrated single-cell RNA sequencing data from fresh human CRPC cases. This comprehensive approach allowed us to identify distinct NEPC subpopulations and their respective lineage with high confidence.

**Key Findings and Limitations:** We uncovered N-Myc and REST as key transcription factors driving distinct neuroendocrine subtypes among 5,797 neuroendocrine-like epithelial cells in CRPC: a REST-dependent subtype (NE I), an N-Myc-dependent subtype (NE II), and a combined N-Myc/REST subtype (NE I+II). These subtypes were validated using multiplex immunofluorescence staining. Trajectory analysis of single-cell RNA sequencing data, along with multi-omics time course analysis of publicly available transcriptomic data recapitulated N-Myc and REST lineages. Additionally, we observed PSMA loss in N-Myc lineage NEPC and identified STMN1 as a biomarker for PSMA-negative subtype. We validated the prognostic value of STMN1 using the TCGA dataset and 60 in-house CRPC tissues. Given that surgery is rarely performed in advanced CRPC, leading to limited sample availability, further validation in larger cohorts is needed.

**Conclusions and Clinical Implications:** Adeno-to-neuroendocrine lineage transition in prostate cancer leads to resistance to new therapies. The lethal NEPC phenotype should be revealed earlier in the disease course of patients with CRPC, providing crucial clues for personalized precision medicine.

## 1. Introduction

Prostate cancer (PCa) is predominantly a hormone-sensitive adenocarcinoma expressing androgen receptor (AR). Androgen deprivation therapy (ADT) initially controls recurrent and metastatic PCa by inhibiting AR signaling, but with prolonged treatment, it develops castration-resistant prostate cancer (CRPC). The prevalence of CRPC led to the development of second-generation anti-androgen receptor (anti-AR) therapies [1, 2]. These treatments, while effective initially [3, 4], can induce lineage plasticity in CRPC, leading to approximately 20% of CRPC patients transitioning to highly aggressive AR-negative neuroendocrine carcinoma (NEPC) [5]. This histological variant PCa is more likely to have visceral metastases and a worse prognosis [6]. Epigenetic reprogramming mediates the transdifferentiation of prostate epithelial cell into neuroendocrine cells, and this often occur in the context of genetic mutations, such as TP53, RB1, and PTEN loss [7, 8]. N-Myc, a master transcription factor driving lineage plasticity, is overexpressed in NEPC and cooperates with FOXA1 or MUC1-C to activate EZH2, SOX2, BRN2/POU3F2, and NKX2-1/TTF1 [9-13]. This cooperation induces neuroendocrine-related genes, highlighting N-Myc as a key driver of neuroendocrine differentiation. Conversely, the loss of RE-1 silencing transcription factor (REST) drives neuroendocrine differentiation in PCa cells by de-repressing neuroendocrine-related genes, underscoring REST as another critical driver of neuroendocrine differentiation [14, 15]. Comprehensive bulk RNA sequencing (RNA-seq) of metastatic castration-resistant prostate cancer (mCRPC) reveals the presence of two distinct neuroendocrine gene profiles, with one highlighting the central role of REST (NEURO I) and the other possessing multiple transcription factors (NEURO II) [16]. Single-cell RNA sequencing (scRNA-seq) data across various histological subtypes of PCa specimens validated the existence of neuroendocrine subtypes [17-19].

In 2022, WHO defined CRPC with partial or complete neuroendocrine differentiation as treatment-induced neuroendocrine prostatic carcinoma (t-NEPC) [20]. In clinical practice, NEPCs are defined by positive immunohistochemical (IHC) staining of neuroendocrine markers, such as synaptophysin (SYP) or chromogranin A (CHGA) [21]. However, current histologic and IHC staining features cannot distinguish NEPC subtypes. Molecularly defined subtypes offer promising prognostication power [16, 19].

## 2. Material and method

### 2.1. Human sample collection

From 2020 to 2022, a total of four fresh metastatic CRPC (mCRPC) and one fresh hormone-sensitive prostate cancer (HSPC) tissue were collected at Taipei Veterans General Hospital (TVGH), including three mCRPCs to lymph node obtained through inguinal lymph node biopsy or pelvic lymph node dissection, and one mCRPC to adrenal with adenocarcinoma histology obtained through laparoscopic adrenalectomy. The primary HSPC (Gleason score 3 + 4) was obtained through robotic-assisted laparoscopic prostatectomy (RALP). Half of the fresh tissue was allocated for formalin-fixed paraffin-embedded (FFPE) diagnosis, and the other half was used to prepare single cells. Informed consent was obtained from the patient before sampling, and experiments with human subjects were approved by the Institutional Review Board (IRB 2020-01-015BC) of Taipei Veterans General Hospital (TVGH).

### 2.2 Single-cell RNA sequencing (scRNA-seq) and data processing

The surgically removed PCa tissues were immersed in MACS Tissue Storage Solution (Miltenyi Biotec, San Diego, CA, USA. cat. no. 130-100-008) and immediately transported to the laboratory on ice. Single cells were prepared using Human Tumor Dissociation Kit (Miltenyi Biotec, San Diego, CA, USA. cat. no. 130-095-929). The tissue block was immersed in 5 ml of EpiGRO™ Human Prostate Epithelial Basal Medium (Millipore, Billerica, MA, USA. cat. no. SCMP-BM) in a sterile Petri dish and minced into small cubes (∼1 mm). The cubes were transferred to a gentleMACS C tube (Miltenyi Biotec, San Diego, CA, USA. cat. no. 130-093-237) containing digestive enzymes diluted in 5 ml of EpiGRO™ Human Prostate Basal Medium. The single-cell suspension was made using gentleMACS Octo Dissociator (program h_TDK_2, Miltenyi Biotec, San Diego, CA, USA. cat. no. 130-096-427). After run completion, 5 ml of EpiGRO Human Prostate Basal Medium was added to the gentleMACS C tube. Then the single-cell suspension was passed through a 70-μm cell strainer (FALCON, Glendale, AZ, USA, cat. no. 352350). Viable cells were enriched by Ficoll density gradient centrifugation (1,000 x g at room temperature for 15 min). The cell number and viability were analyzed using Countess II FL Automated Cell Counter (Invitrogen, Carlsbad, CA, USA. cat. no. AMQAF1000). The single-cell suspension with viability >90% was centrifuged (300 x g at 4°C for 7 min) and resuspended at a density of 1,000 cells/μl in EpiGRO™ Human Prostate Basal Medium. The suspension of 8,000-10,000 cells was encapsulated into single droplets using Single-Cell-A Chip (10x Genomics, San Francisco, CA, USA. cat. no. PN-1000121) for target capture. The libraries were prepared using Chromium Next GEM Single Cell 3□Reagent Kits v3.1 (10x Genomics), according to the manufacturer’s protocol. The library was sequenced on a NovaSeq 6000 (Illumina, San Diego, CA, USA. cat. no. 20012850) as 150bp pair-end reads with a depth of ∼150 M reads at National Genomics Center for Clinical and Biotechnological Applications, Cancer Progression Research Center (National Yang Ming Chiao Tung University).

Sequencing data from each library were processed using the Cell Ranger (10x Genomics, version 7.0.0) analysis pipeline for demultiplex, barcode processing, and converted to FASTQ format, followed by alignment to the human reference genome (GRCh38) and acquire alignment matrix file. Reads that passed QC were processed through CellRanger count to obtain unique molecular identifiers (UMIs) matrices. Barcodes with low unique molecular identifier (UMI) counts, low feature counts, and high percentages of mitochondrial gene expression were excluded. Doublets were detected and removed using DoubletFinder [22]. The scRNA-seq data from four CRPC samples in GSE137829, including three with NEPC characteristics (GSE#2, GSE#5, and GSE#6), and one identified as CRPC (GSE#4), were processed in accordance with the methodologies outlined in the corresponding publication [18]. Further analysis was conducted using Partek Flow (Partek, Cat. No. 4485102), including normalization with log2(CPM+1), noise reduction by excluding genes expressed in fewer than 0.1% of cells, batch effect removal using the Seurat v3 integration method [23], and dimensionality reduction via principal component analysis.

### 2.3 Cell type identification and PCa cell classification

Using principal component 15 and a resolution of 0.5, we identified 18 cell clusters through graph-based clustering for non-linear dimension reduction and visualized using a t-SNE plot. The cell types of each cluster were annotated based on canonical markers from Song *et al*. Clusters exhibiting high expression of epithelial markers, particularly EPCAM, were classified as PCa epithelial cells and selected for further analysis.

To explore the heterogeneity of PCa cells, epithelial cells were combined and re-clustered using principal component 14 and a resolution of 0.5. We assessed each cluster for their expression of commonly recognized RNA expression signature for prostate adenocarcinoma (Liu’s PCa score [24] & Tomlins’ PCa score [25]), club cells [26], androgen response [27], neuroendocrine (NEURO I score and NEURO II score [16]), Rb inactivation [28], TP53 depletion [29], TMPRSS2-ERG fusion [30], and Wnt/β-catenin [27]. Violin plots were constructed using the score of RNA signature.

To investigate the regulation of key neuroendocrine-related transcription factors in different neuroendocrine clusters, the NRSF_01 and NMYC_01 gene sets were downloaded from MsigDB [27, 31] and used for gene set enrichment analysis (GSEA) [32, 33]. GSEA was performed to analyze the enrichment in NE I (Epi-C11), NE II (Epi-C6), and NE I+II (Epi-C5 and Epi-C7) relative to all other clusters using Partek Flow.

### 2.4 Single-cell trajectory analysis and biomarker prediction

To analyze NEPC progression, single-cell pseudotime trajectory of epithelial cells was constructed by Monocle 2 [34] in Partek Flow. Using Reversed Graph Embedding, each epithelial cell was projected onto a manifold in a low-dimensional space, which ordered the cells into a trajectory and identified any branch points corresponding to cell fate decisions. Violin plots were constructed using the AR, NEURO I, and NEURO II scores. The Hallmark AR, NRSF_01, and NMYC_01 gene sets were used for GSEA enrichment analysis of state 1 and state 2 relative to all other states.

To identify potential biomarkers predicting the prognosis of CRPC with neuroendocrine features at an early stage, we first performed correlation analysis on highly expressed (expression level >5) biomarkers in each NEPC evolutionary trajectory state (*p*<0.05) and calculated their association with NEURO scores. We distinguished 235 biomarker genes in state 1 that correlate with the NEURO I+II score, 68 biomarker genes in state 2 that correlate with the NEURO II score, and three biomarker genes in state 5 that correlate with the NEURO I score. Among these, 47 of the 235 biomarkers in state 1 and 9 of the 68 biomarkers in state 2 predicted poor prognosis of PCa in the TCGA dataset.

### 2.5 Verification using public animal and clinical databases

We employed scRNA-seq and bulk RNA-seq data from the PARCB Time Course Multiomics Explorer website (https://systems.crump.ucla.edu/transdiff/index.php) [35] to evaluate the expression levels of REST, MYCN, and our potential NEPC markers in NEPC progression. The NEPC dataset from the Beltran Cohort was downloaded from cBioPortal and used to perform receiver operating characteristic (ROC) curve analysis of our potential NEPC markers using GraphPad Prism (version 10.1.1) [36]. Youden’s index was used for the identification of the optimum cut-off point for STMN1. The ROC curve of STMN1 indicated that the best cutoff was 0.1833 in predicting NEPC. Prostate cancer cohorts from MSKCC [37], available on the camcAPP website (http://bioinformatics.cruk.cam.ac.uk/apps/camcAPP/) [37], were used to perform survival analysis of our potential NEPC markers using camcAPP. The expression levels of our potential NEPC markers in various PCa cell lines, including two non-cancerous prostates, nine prostate adenocarcinoma, and one prostate small cell carcinoma (NCI-H660), were analyzed by using the DepMap portal (https://depmap.org/portal).

### 2.6. Multiplex immunofluorescence and immunohistochemistry staining

Multiplex IF staining of FFPE tissue was performed by Opal Multiplex Immunostaining kit Opal ^™^ 6-PLEX MANUAL DETECTION KIT (Akoya Biosciences, Marlborough, MA, USA. cat. no. NEL 861001KT) following the manufacturer’s instructions. FFPE sections were deparaffinized in xylene, rehydrated through graded ethanol series (100%, 95%, and 70%), and subjected to antigen retrieval in an autoclave at 121°C and 1 atm for 10 minutes. This was done using either pH 6.0 (AR6 buffer, cat. no. S1699) or pH 9.0 (AR9 buffer, cat. no. S2367) Target Retrieval Solution (Agilent, Santa Clara, CA, USA). Cytokeratin 8 [C-43] [Abcam, ab2530], synaptophysin [Abcam, ab32127], N-Myc [Cell Signaling Technology, #51705], REST [Invitrogen, MA5-24606], antibodies were sequentially applied, followed by incubation with Opal Polymer HRP Ms+Rb and Opal fluorophore-conjugated tyramide signal amplification. After each signal amplification step, slides were heat-treated in an autoclave. Nuclei were stained with DAPI after all antigens had been labeled. Slides were scanned to acquire multispectral images using the Vectra® Polaris™ Automated Quantitative Pathology Imaging System (PerkinElmer, Waltham, MA, USA. cat. no. CLS143455).

For Immunohistochemical (IHC) staining, FFPE sections were deparaffinized by EZ prep (Ventana Medical Systems, Inc., Tucson, AZ, USA) and stained with specific antibodies of Stathmin [Cell Signaling Technology, #13655] at room temperature for 32 minutes using the automated Ventana Benchmark XT (Ventana Medical Systems, Inc., Tucson, AZ, USA). Slides were visualized using Ultraview DAB Detection Kit (Ventana Medical Systems, Inc., Tucson, AZ, USA) in the presence of hematoxylin counterstaining following the manufacturer’s protocol. The staining histology was scanned in five fields for each sample, and H-score was calculated based on a four-tier scale: negative (0), weak (1), intermediate (2), and strong (3). Kaplan-Meier estimates overall survival (OS). The OS was defined as the time from tissue biopsy to death from any cause.

## 3. Result

### 3.1. Intrinsic NEPC subpopulations in CRPC

To characterize intratumor neuroendocrine heterogeneity, we collected fresh PCa tissues, including four mCRPC and one prostatectomy hormone-sensitive prostate cancer (HSPC) (Table 1). The neuroendocrine feature was confirmed in three of the four mCRPC cases and one HSPC (Fig. 1A). Enriched viable cells from each sample were subjected to droplet-based single-cell RNA sequencing (scRNA-seq). To increase sample size, we downloaded four CRPC samples from GSE137829, of which three exhibited NEPC characteristics and one was identified as CRPC. After quality control, we obtained 29,805 single cells and identified 18 distinct cell clusters (Supplementary Fig. S1A and S1B). Cell types for each cluster were defined using canonical markers specific to each cell type (Fig. 1B, 1C, and Supplementary Fig. S1C and S1D). Consistent with PCa being considered a cold tumor, most PCa samples exhibited low to no immune cell populations, except for GSE#5, which had abundant immune cells, and PCa-2, PCa-3, and GSE#6, which had some immune cells (Fig. 1D).

**Table 1.** Clinical characteristics.

| Patient ID | PCa-1 | PCa-2 | PCa-3 | PCa-4 | PCa-5 |
| --- | --- | --- | --- | --- | --- |
| Age at sequencing | 59 | 61 | 60 | 65 | 69 |
| Sequencing date | 2020/3/17 | 2020/10/27 | 2022/3/31 | 2022/6/10 | 2022/6/21 |
| PSA level at sequencing | 745 | 20.8 | 7.29 | 1886 | 0.14 |
| Biopsy site | lymph node | Adrenal | Prostate | lymph node | lymph node |
| TMN stage | cTxcN0M1a+b | cTxcN0M1a+b+c | cT3acN0M0 | cT4cN1M1a+b | cT4cN1M0 |
| Gleason score | None | 5+5 | 3+4 | None | 4+5 |
| Disease status when sequencing | CRPC + NEPC | CRPC | HSPC | CRPC | Mixed small cell<br>NEPC + adeno G4+5 |
| Drug treatment after first diagnosis | ADT + RT | ADT + RT | ADT | ADT | Etoposide + Cisplatin |
| Second line treatment | Doxcetaxel | Doxcetaxel | None | None | None |
| Death date | 2021/10/20 | 2021/2/1 | Alive | 2023/6/21 | 2023/2/25 |

CRPC: Castration-resistant prostate cancer; NEPC: Neuroendocrine prostate cancer; ADT: Androgen deprivation therapy; RT: Radiation therapy

**Fig. 1.**
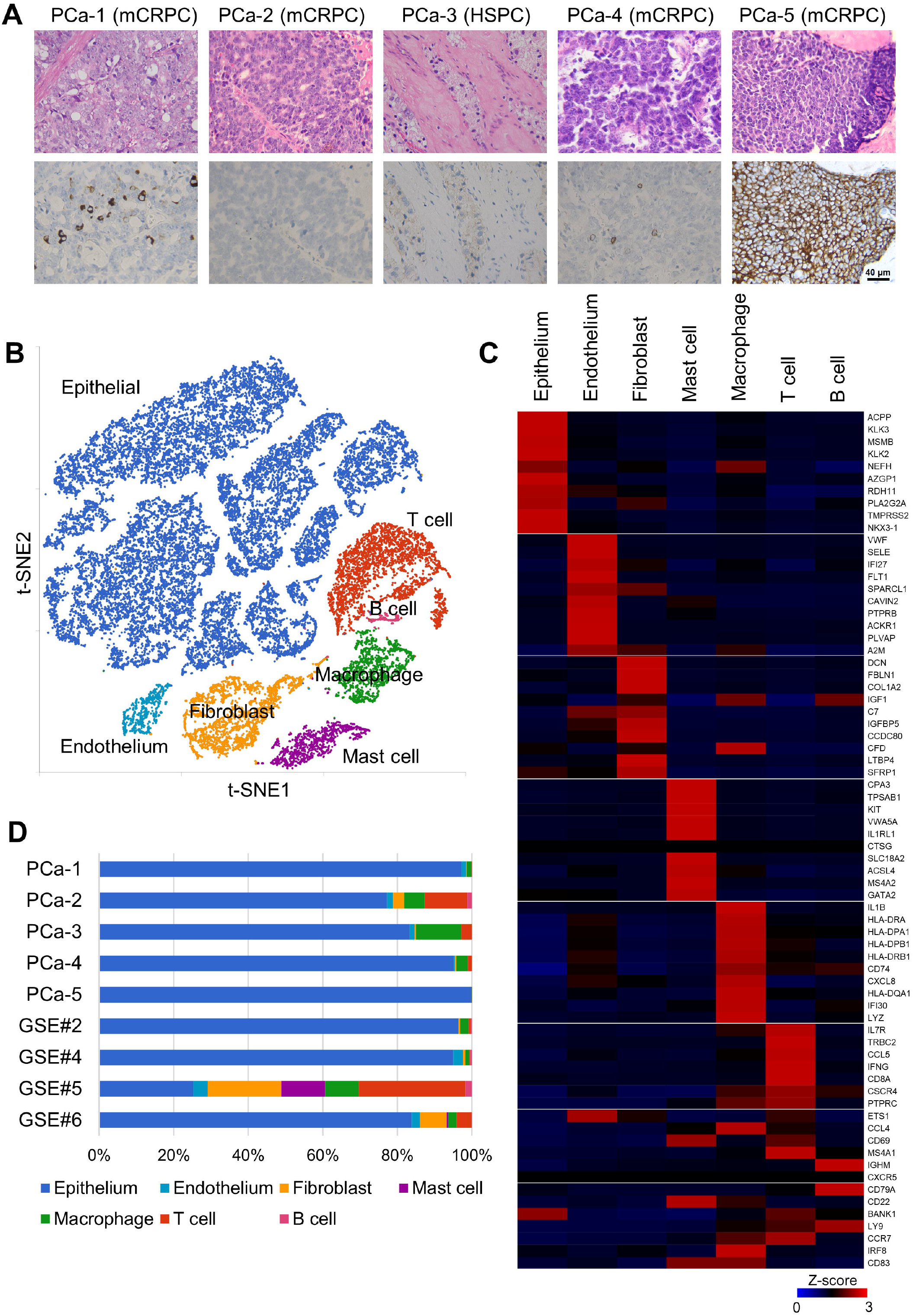
Prostate epithelial cells in CRPC and HSPC were identified in scRNA-seq. **(A)** Representative histological (upper panel) and IHC (lower panel) images of the four mCRPC (PCa-1, PCa-2, PCa-4, and PCa-5) and an HSPC (PCa-3). IHC observation of neuroendocrine marker SYP. **(B)** t-SNE plot of all major cell types that passed quality control, derived from four mCRPC (PCa-1, PCa-2, PCa-4, and PCa-5) and one HSPC (PCa-3) (n=12,696), along with four locally recurrent CRPC samples (GSE#2, GSE#4, GSE#5, GSE#6) from GSE137829 (n=17,109). Labeled by cell type. **(C)** Heatmap demonstrates marker genes expressed in each cell type. **(D)** Distribution of cells isolated from eight CRPC and one HSPC. mCRPC = metastatic CRPC; HSPC = hormone-sensitive prostate cancer; PCa = prostate cancer; scRNA-seq = single-cell RNA sequencing; SYP = synaptophysin; IHC = immunohistochemical; t-SNE = T-distributed stochastic neighbor embedding.

After collecting epithelial clusters, we performed clustering analysis again and identified 13 distinct PCa epithelial clusters (Epi-C) (Fig. 2A). We then conducted Seurat AddModuleScore analysis on all Epi-Cs using various gene set signatures (Supplementary Table S1 and Supplementary Fig. S2A and S2B) and defined multiple epithelial subpopulations with high PCa and androgen response scores as adenocarcinoma (Fig. 2B, gray). We also found a subpopulation with high club cell score [38] (Fig. 2B, purple), as well as a subpopulation with low androgen response score without gain of NE markers (AR^Low^/NE^Low^), known as double-negative prostate cancer (DNPC) cells (Fig. 2B, green). Importantly, we identified four epithelial cell clusters with high neuroendocrine scores: one with high NEURO I score (Fig. 2B, blue), one with high NEURO II score (Fig. 2B, red), and two with high scores in both NEURO I and NEURO II (Fig. 2B, yellow). Thus, we classified them as NE I, NE II, and NE I+II clusters (Fig. 2B).

**Fig. 2.**
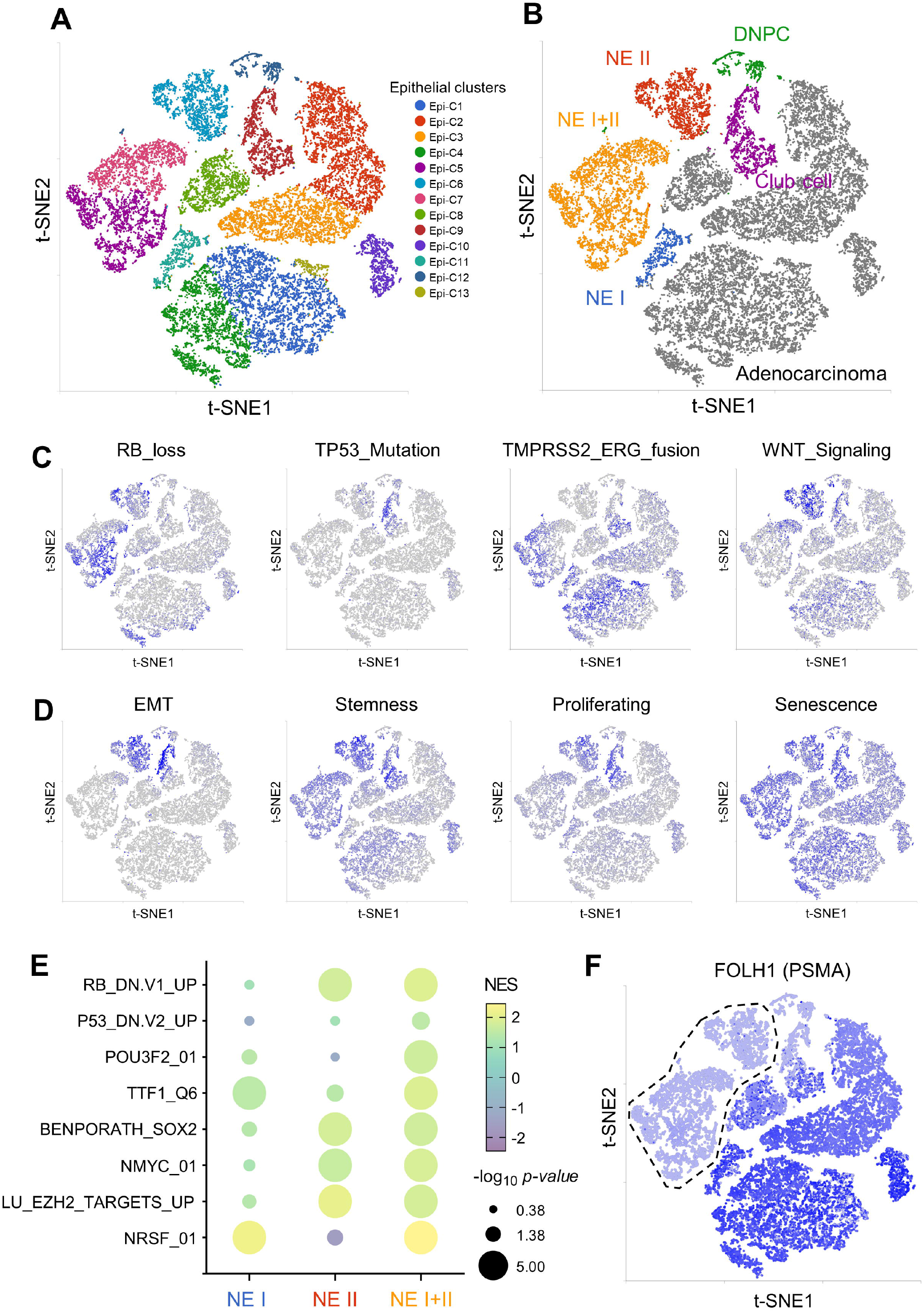
Luminal cells display distinct neuroendocrine phenotypes and regulated by unique transcription factors. **(A)** t-SNE plot of all epithelial cells (n=21,434) colored by the 13 clusters. **(B)** t-SNE view of clusters colored by lineage subtypes. Cell clusters were classified as adenocarcinoma, DNPC (AR^Low^/NE^Low^), club cell, NE I with a high NEURO I score, NE II with a high NEURO II score, and NE I+II with a high NEURO I+II score. **(C-D)** t-SNE plot of cellular pathway signaling (C) cellular gene set signature scores (D). The expression level is presented by color, from no expression (gray) to high-level expression (blue). **(E)** GSEA analysis of DEGs in NE I, NE II, and NE I+II versus all other clusters based on the gene expression profiles of RB_DN.V1_UP, P53_DN.V2_UP, POU3F2_01, TTF1_Q6, BENPORATH_SOX2_TARGETS, NMYC_01, LU_EZH2_TARGETS_UP, and NRSF_01. **(F)** t-SNE plot view of PSMA. DNPC = AR and NE double-negative prostate cancer. AR = Androgen response; NE = neuroendocrine; GSEA = Gene Set Enrichment Analysis; DEGs = differentially expressed genes. NRSF = neuron-restrictive silencer factor.

### 3.2. NEPC heterogeneity in relation to poor prognosis and disease progression

To identify functional differences among neuroendocrine cell clusters, we evaluated RNA expression signatures associated with PCa, including RB inactivation, TP53 depletion, TMPRSS2-ERG fusion, and Wnt/β-catenin signaling [39] (Supplementary Table S1) in all Epi-Cs using gene list from MSigDB (Fig. 2C). We observed higher RB inactivation in the NE II and NE I+II clusters, while the NE I cluster exhibited a pattern similar to all other Epi-Cs. The similarity of most signatures across Epi-Cs supports the notion that NEPC develops from prostate adenocarcinoma, while unique pathways may drive NEPC lineage plasticity. Additionally, Wnt/β-catenin signaling was more active in the NE II and club cluster (Fig. 2C). Wnt/β-catenin pathway is known to regulate pluripotency, proliferation, EMT/migration/invasion/metastasis, stemness, and therapy resistance [40]. Aligned with this, we found upregulated proliferation, EMT, and stemness signatures in NE II and club clusters, with elevated stemness signatures in NE I+II cluster (Fig. 2D). Since EMT promotes stemness [41], these findings suggested correlated lineage plasticity in NE II and NE I+II. This is consistent with advanced NEPC shared stemness and neuronal signatures [42, 43].

### 3.3. Critical transcription regulators modulate PCa progression into different NEPC subtypes

Transcription factors drive tumor lineage plasticity. GSEA analysis showed that NE I was associated with significantly increased REST-repressed genes, indicating a loss of REST activity (Fig. 2E, left). NE II was associated with the loss of Rb and a significant increase in N-Myc-targeted genes (Fig. 2E, middle). Moreover, NE I+II was associated with the loss of Rb and increased N-Myc-targeted and REST-repressed genes, suggesting potential lineage plasticity in these cells (Fig. 2E, right). We confirmed the loss of REST and overexpression of N-Myc in neuroendocrine-positive (SYP^+^) cells in PCa-1 and PCa-5, respectively, using multiplex immunofluorescence (IF) staining (Supplementary Fig. S2C), suggesting the presence of NEPC-REST and NEPC-N-Myc lineage in CRPC.

### 3.4. Heterogeneity of NEPC and its implications for prostate-specific membrane antigen (PSMA)-targeted therapies

The PSMA-targeted radionuclide ^177^Lu-PSMA-617, a radiopharmaceutical, was approved by the US Food and Drug Administration (FDA) in 2022 for men with CRPC. It is indicated for PSMA-positive mCRPC patients previously treated with anti-AR signaling therapies and taxane-based chemotherapy [44]. PSMA, an AR-repressed transmembrane glycoprotein, is upregulated in prostate adenocarcinoma by 100-to 1000-fold and tends to increase as tumors progress to CRPC [45]. PSMA cistrome analysis across PCa progression also identifies PSMA-positive NEPC tumors [46]. Despite these findings, PSMA expression is often suppressed in AR-negative advanced PCa, presenting a challenge for patient management and highlighting the need for synergistic combination therapies. Therefore, we further color-coded Epi-Cs with FOLH1 (PSMA) (Fig. 2F). Interestingly, we found that NE I cluster expressed PSMA at levels comparable to CRPC adenocarcinoma cells, while clusters with NE II features (NE II and NE I+II) exhibited low to no PSMA expression. These findings align with clinical observations, indicating the presence of both PSMA-positive and -negative NEPC tumors. More importantly, our results suggest that patients with high NE I feature could be effectively managed with ^177^Lu-PSMA-617. In contrast, patients with high NE II features may require combination therapy to induce PSMA expression, making them responsive to ^177^Lu-PSMA-617.

### 3.5. The trajectory depicted CRPC progression toward distinct NEPC subtypes

Next, we explored correlations between cellular heterogeneity and CRPC progression to NEPC using the Monocle 2 reverse graph embedding approach. we decomposed PCa epithelial cells into a tetrafurcated trajectory, including paths toward the NE II subtype ending with NE I+II cluster in state 1 and DNPC in state 2 (Fig. 3A). We also identified two adenocarcinoma trajectories: one with (state 5) and one without (state 4) the NE I cluster, with club cells positioned at interphase (state 3) (Fig. 3A, right). NE I cells resided mainly in a single CRPC adenocarcinoma trajectory (state 5) indicating their closer relationship to CRPC adenocarcinoma. NE II cells were distributed across state 1 (leading to NE I+II clusters) and state 2 (leading to AR^Low^/NE^Low^ DNPC), suggesting NE II cells may acquire NE I features or transition to DNPC, potentially increasing CRPC lineage plasticity under hormone therapy. By calculating androgen response, NEURO I, and NEURO II scores, we validated the evolutionary trajectory state 1 carrying both NEURO I and NEURO II signatures (Fig. 3B). GSEA analysis further confirmed that state 1 was negatively associated with androgen response and positively related to N-Myc-targeted and REST-repressed genes (Fig. 3C). Importantly, color-coded trajectories with FOLH1 (PSMA) revealed that states 1 and 2 exhibited low to no PSMA expression (Fig. 3D and 3E). Consequently, we defined NE II and NE I+II clusters as PSMA-negative NEPC. Additionally, PSMA was also lost in DNPC (Fig. 3D and 3E). Since NE II spans states 1 and 2, we hypothesize that N-Myc may play a critical role in PSMA suppression.

**Fig. 3.**
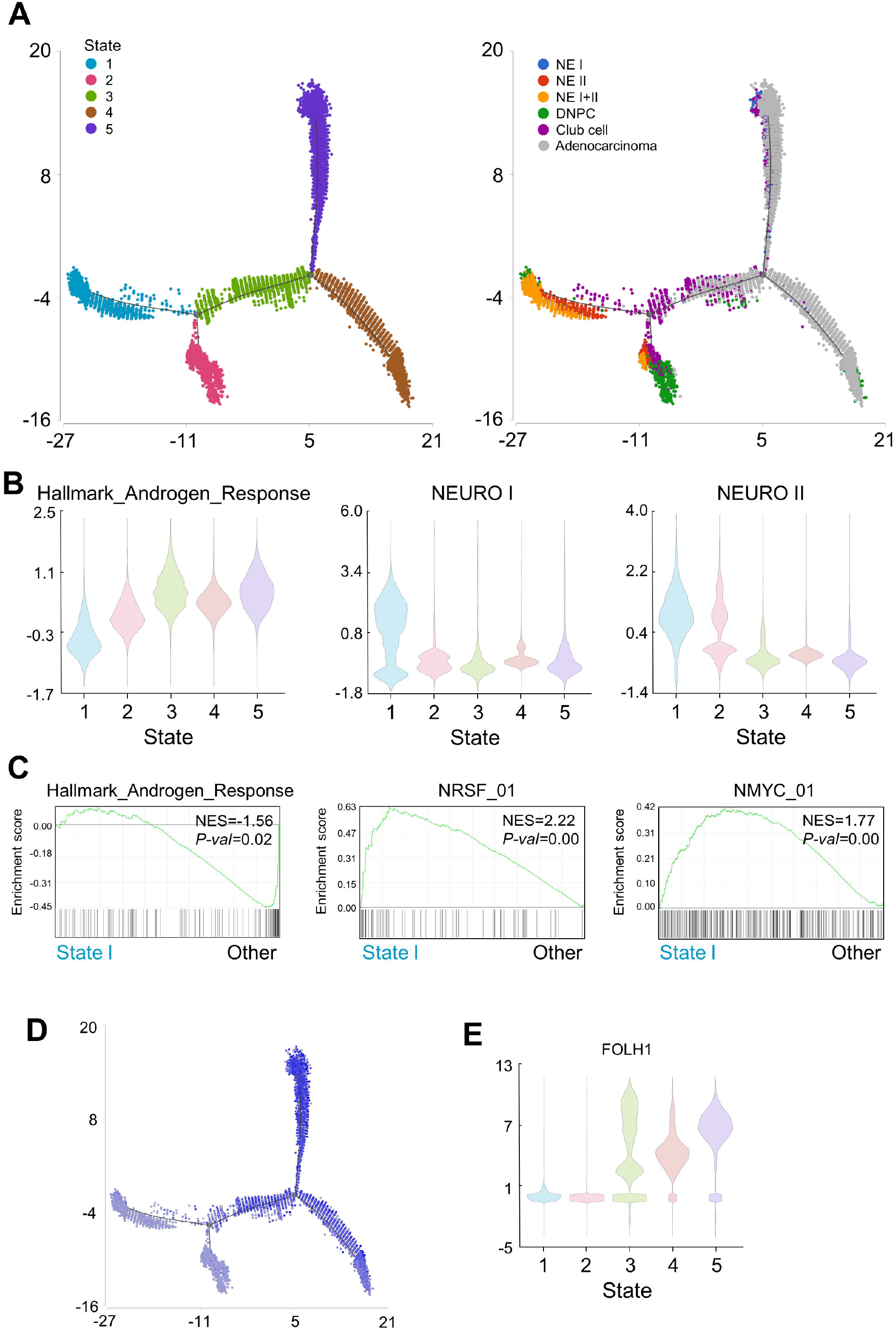
Trajectory analysis of luminal cells reveals cell states representing NE I+II features. **(A)** Trajectory plot of luminal cells, color-coded by cell states (left panel) and by cell types identified in Fig. 2B (right panel). **(B)** Violin plots visualizing AR, NEURO I, and NEURO II scores among luminal cells within each cell state in the trajectory. **(C)** GSEA analysis of biomarkers in state 1 versus all other states, based on gene expression profiles of AR Hallmark, NRSF_01, and NMYC_01. **(D)** Trajectory plot of luminal cells, color-coded by PSMA expression. **(E)** Violin plots visualizing PSMA expression in each cell state in the trajectory.

### 3.6. Highly plastic NEPC cells were observed in an independent dataset

To validate the association of NEPC subtypes with REST and N-Myc, we reanalyzed REST and MYCN expression in the multi-omics time course dataset of PARCB, a pan-small cell neuroendocrine cancer mouse model [35]. Our analysis revealed a temporal up-regulation of MYCN and down-regulation of REST in the PARCB time course (Supplementary Fig. S3A). Additionally, the PARCB transformation trajectory showed MYCN up-regulation in transitional tumors (HC4) and late tumors (class 1; HC5) at one endpoint of the trajectory (Supplementary Fig. S3B, left), whereas REST was down-regulated in another set of late tumors (class 2; HC6) at the opposite endpoint of the trajectory (Supplementary Fig. S3B, right). These results supported our discovery that REST and N-Myc are the key driving transcription regulators that modulate differential neuroendocrine differentiations.

### 3.7. Identification of STMN1 as a novel diagnostic marker for PSMA-negative NEPC

To identify potential biomarkers for the diagnosis and prognosis of PSMA-negative NEPC at an early stage, we conducted correlation analysis of NEURO scores, disease-free survival analysis using TCGA dataset, and receiver operating characteristic (ROC) curve analysis using Beltran’s NEPC dataset [36] on highly expressed biomarkers (expression level >5, *p*<0.05) in state 1 that correlate with the NEURO I+II score (Supplementary Table S2). This analysis identified seven biomarkers from state 1 with poor disease-free survival (log-rank *p*<0.05) (Supplementary Table S2) and a high area under curve (AUC > 0.8) (Fig. 4A). The prognostic value of STMN1, UBE2S, PTTG1, HES6, and KALRN was validated using MSKCC dataset [37] (Fig. 4B).

**Fig. 4.**
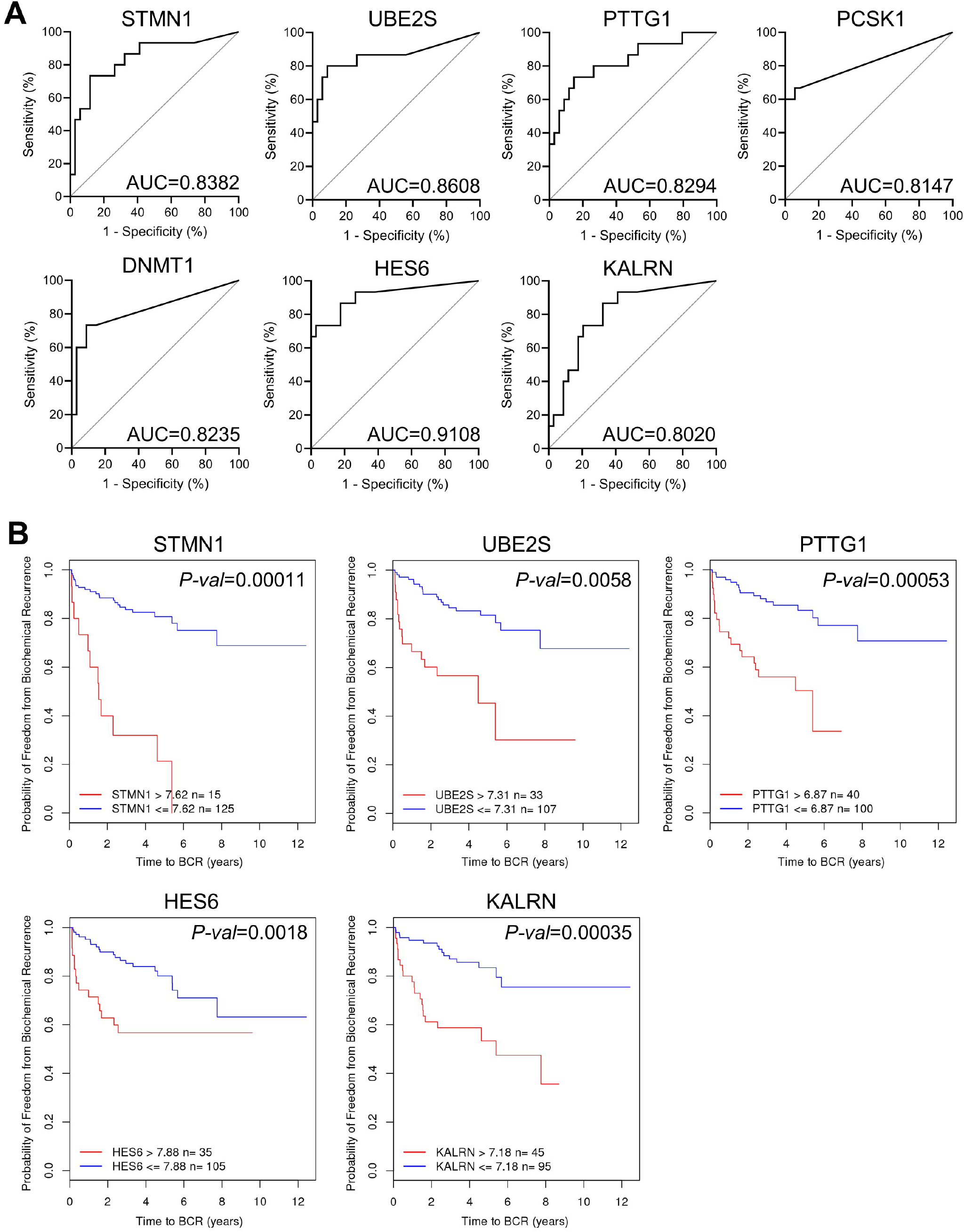
Clinical relevance of NEURO I+II correlated poor prognostic biomarkers in state 1 for NEPC diagnosis and prognosis. **(A)** ROC curve of seven biomarker genes from state 1 in distinguishing NEPC from PCa using Beltran’s NEPC dataset. **(B)** Kaplan-Meier plots showing the biochemical recurrence-free rate in PCa for five of the seven biomarkers: STMN1, UBE2S, PTTG1, HES6, and KALRN from the MSKCC dataset. ROC = receiver operating characteristic.

Using the PARCB model, we observed an increase in STMN1, UBE2S, HES6, and KALRN in early tumors (HC3) and significantly higher levels in intermediate (HC4) and late tumors (HC5 and HC6) compared with prostate basal (HC1) and organoid (HC2) (Supplementary Fig. S4A). Notably, STMN1 and UBE2S were significantly higher in N-Myc up-regulated late tumors (HC4 and HC5) compared with REST down-regulated late tumors (HC6) (Supplementary Fig. S2B and S4A). We further validated the association of these biomarkers with N-Myc expression in PCa cell lines. STMN1 was significantly higher in human NEPC cells (Supplementary Fig. S4B), indicating STMN1 as a valuable biomarker for NEPC-N-Myc in both preclinical models and clinical samples. To further validate STMN1 as a biomarker for CRPC prognosis, we performed immunostaining on 60 CRPC tissues and analyzed the immunostaining with H-score (Fig. 5A). CRPC with high STMN1 expression (H-score >60) showed significantly lower overall survival (Fig. 5B). Using Beltran’s dataset [31], we confirmed a negative correlation between PSMA and STMN1 in NEPC (Fig. 5C) and validated low PSMA expression in STMN1-high CRPC (Fig. 5D). The identification of STMN1 as a potential diagnostic marker for PSMA-negative NEPC will enhance precision medicine approaches to PSMA theranostics.

**Fig. 5.**
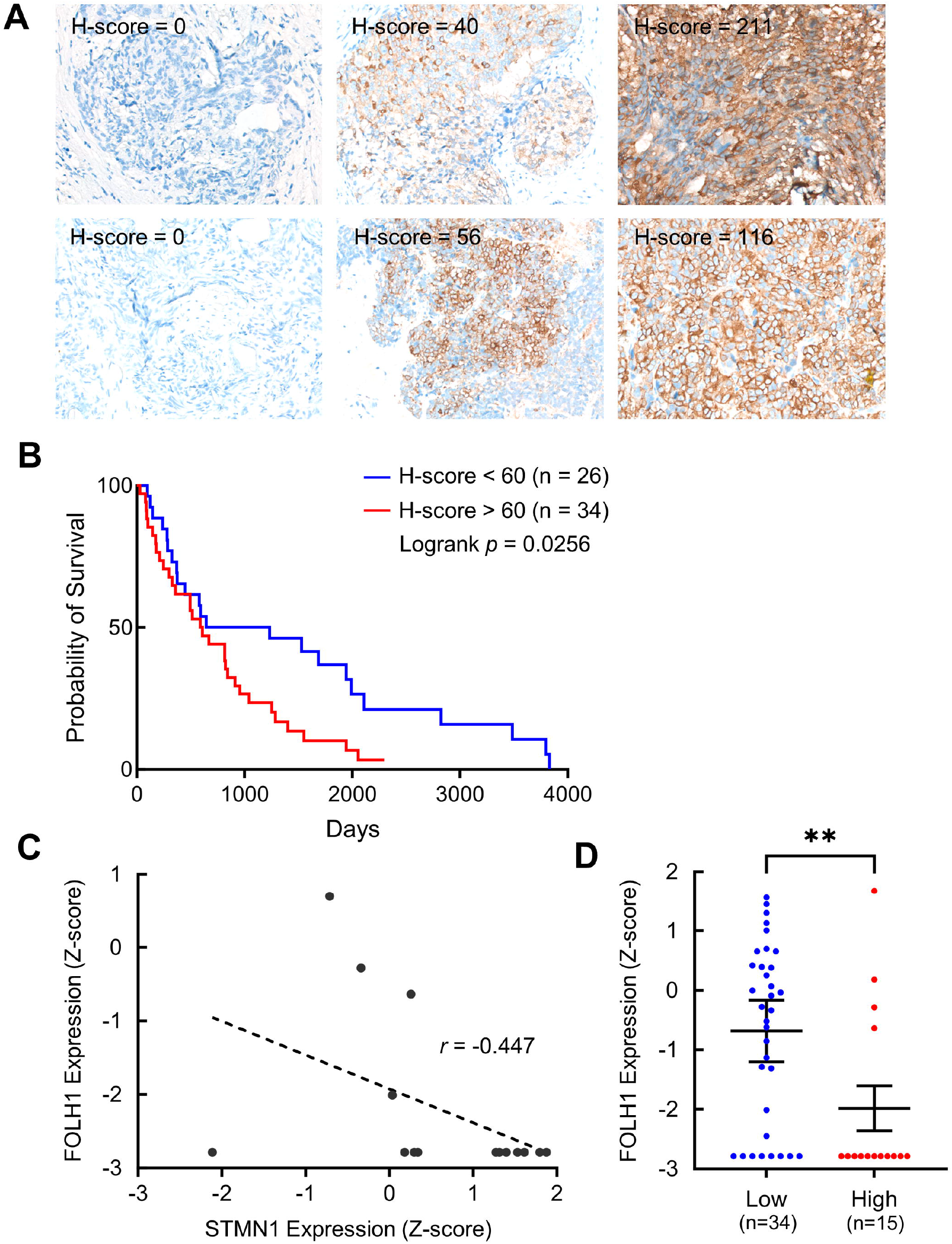
Clinical relevance of STMN1 expression in predicting the survival of CRPC patients. **(A)** Representative immunostaining images of STMN1 in CPRC with low (H-score <60) and high (H-score >60) expression. Positive staining with anti-STMN1 antibody is presented in brown. The cell nucleus was stained with hematoxylin and presented in blue. **(B)** Kaplan-Meier plots predicting poor survival of CRPC patients based on the expression levels of STMN1. **(C)** FOLH1 (PSMA) levels were inversely correlated with STMN1 in NEPC samples in Beltran’s NEPC dataset. **(D)** FOLH1 (PSMA) levels were compared between high and low STMN1 expression groups in Beltran’s NEPC dataset.

## 4. Discussion

We identified three NEPC subtypes and elucidated N-Myc and REST as key transcription factors driving disease progression. N-Myc, a master regulator, directs or indirectly activates NEURO II marker genes [16], including NKX2-1/TTF1 [47], BRN2/POU3F2, and SOX2 [13]. REST inactivation has been shown to transform adenocarcinoma into neuroendocrine carcinoma [48, 49]. Using multi-omics time course analysis of PARCB mouse model implanted with transformed organoids from 10 PCa patients [35], we validated the temporal increase of N-Myc and decrease of REST in divergent NE populations, supporting the notion that N-Myc and REST drive distinct lineage plasticity and neuroendocrine transdifferentiation in t-NEPC. Importantly, we found that N-Myc-regulated NE II and NE I+II NEPC subtypes are mainly PSMA-negative. This aligns with findings that, in the absence of androgens, N-Myc enhances EZH2 recruitment to AR target genes, increasing H3K27me3 levels, and thus suppresses AR-targeted genes [11, 47]. As PSMA is an AR-regulated gene, our results suggest that N-Myc and EZH2-mediated transcriptional reprogramming may underlie PSMA suppression in certain NEPC subtypes. The availability of EZH2 inhibitors, such as the FDA-approved tazemetostat and CPI-1205 in clinical trials [50], enables the combination therapy with ^177^Lu-PSMA-617 for NEPC patients exhibiting low PSMA expression. Additionally, among the NEURO scores-associated biomarkers identified in state 1, STMN1 has been used to predict high-grade lung neuroendocrine tumors [51]. In this study, STMN1 was detected at various levels in CRPC and demonstrated a predictive role in patient survival. This suggests that CRPC patients with high STMN1 expression may harbor pre-existing NE I+II or precursor cells, which could rapidly develop into an aggressive NEPC subtype and become resistant to ^177^Lu-PSMA-617, necessitating alternative interventions. Given the limited CRPC sample size, future prospective studies are needed. Fortunately, current international guidelines recommend biopsy of metastatic lesions in mCRPC. Early identification of STMN1-high and/or PSMA-low lesions may help refine future PSMA theranostics.

## 5. Conclusion

We identified distinct NEPC subtypes with different neuroendocrine signatures in CRPC, underscoring their potential to guide personalized therapy. Notably, NE II and NE I+II subtypes are associated with poor prognosis and diminished PSMA expression, highlighting the need for alternative therapeutic approaches. This study is the first to characterize NEPC subtypes at single-cell resolution, revealing key transcription regulators and their prognostic implications.

---

Fig. S1. - Single-cell transcriptome analysis of HSPC and CRPC. **(A)** t-SNE projection of expression profiles of 29,805 cells isolated from HSPC and CRPC samples. Dots represent each cell, color-coded by cell clusters. **(B)** t-SNE of (A) was color-coded by samples. **(C)** t-SNE view of epithelial marker EPCAM, endothelium marker FLT1, fibroblast marker FBLN1, macrophage marker CD68, T cell marker CD3D, B cell marker CD79A, and mast cell marker MS4A2. **(D)** t-SNE view of prostate luminal cell marker KRT8 and basal cell marker KRT-5. HSPC = hormone-sensitive prostate cancer; CRPC = castration-resistant prostate cancer; t-SNE = T-distributed stochastic neighbor embedding; EPCAM = epithelial cell adhesion molecule; FBLN1 = fibulin-1; CD = cluster of differentiation; MS4A2 = membrane-spanning 4-domains A2. KRT = Keratin.

Fig. S2. - Luminal cells display plastic and neuroendocrine phenotype. **(A)** Feature plots of clusters measured by Liu’s PCa, Tomlins’ PCa, club cell, AR, NEURO I, and NEURO II scores. The expression level was presented by color, from no expression (gray) to high-level expression (blue). **(B)** the t-SNE plot of 21,434 epithelial cells colored by the samples. **(C)** Representative images of multiplex immunofluorescence staining in FFPE tissues collected from PCa-1 and PCa-5 patients staining with anti-SYP (green), anti-REST (yellow), anti-N-Myc (red), and DAPI (blue). (a) A positive signal of SYP but not REST, and N-Myc in PCa-1 indicates the NE I feature. (b) Positive signal of SYP, REST, and N-Myc in part of PCa-5 indicating NE II feature. (c) Positive signal of SYP and N-Myc but not REST in part of PCa-5 indicates the NE I+II feature. GSEA = Gene Set Enrichment Analysis; DEGs = differentially expressed genes; SYP = synaptophysin. AR = androgen response; NE = neuroendocrine; KLK3 = kallikrein-3; PSA = prostate-specific antigen; FOLH1 = folate hydrolase 1; PSMA = prostate-specific membrane antigen.

Fig. S3. - Validation of N-Myc and REST in distinct neuroendocrine differentiation during PCa progression. **(A-B)** Expression of MYCN and REST over the time course of PARCB, including basal, organoid, early, transition, and endpoint (A), and in each HC (B). PARCB = a pan-small cell neuroendocrine cancer model; HC = hierarchical cluster.

Fig. S4. - Validation of the six biomarkers of state 1 in mouse model and human cell lines. **(A)** Box plot of STMN1, UBE2S, PTTG1, HES6, and KALRN expression in each HC. **(B)** Dot plot of the expression of STMN1, UBE2S, PTTG1, HES6, and KALRN in correlation with MYCN.

