## Supplementary figures and images for "Distinct prostate cancer neuroendocrine subtypes predict prognosis and guide personalized prostate-specific membrane antigen-targeted therapy"

### Supplementary Figure

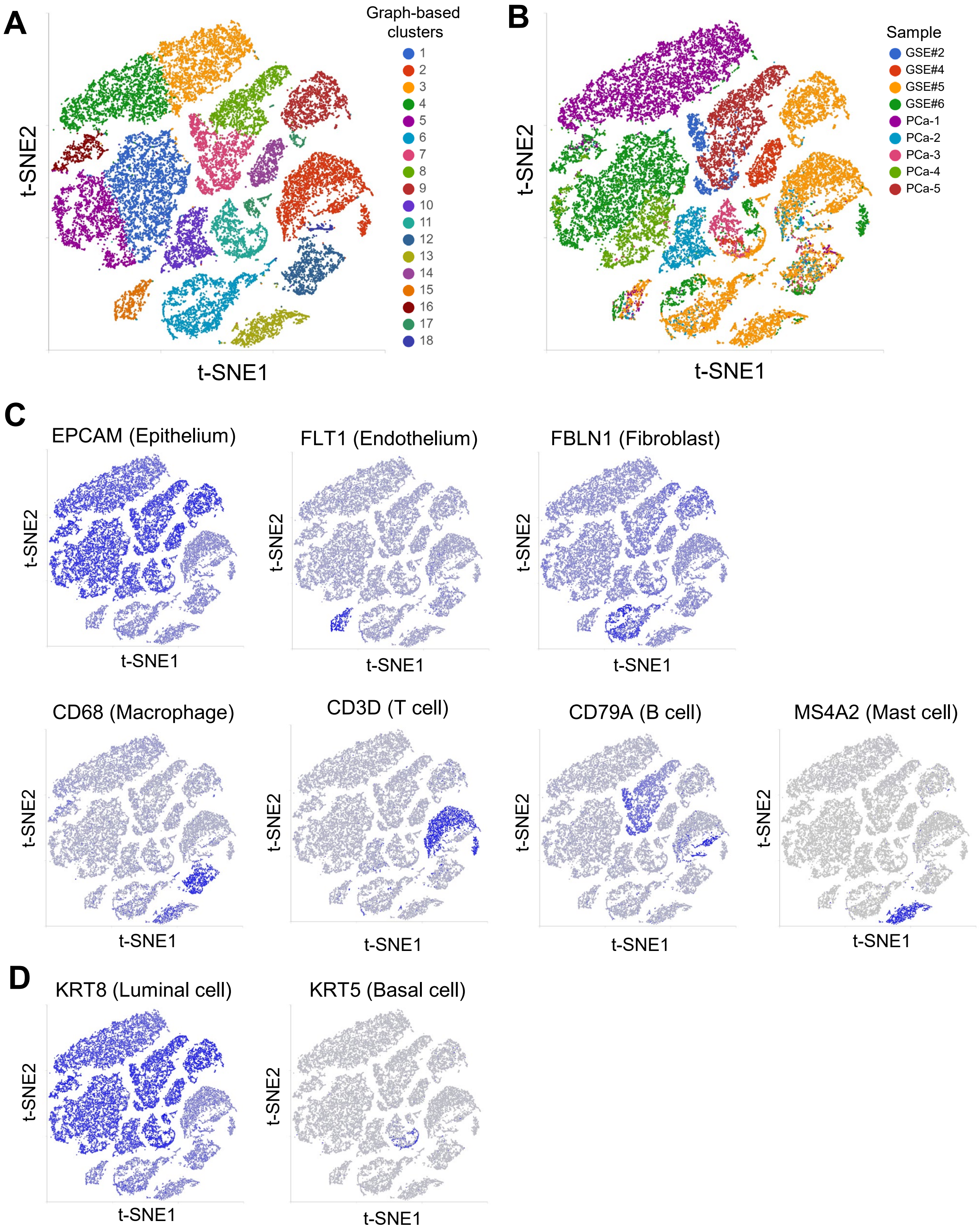

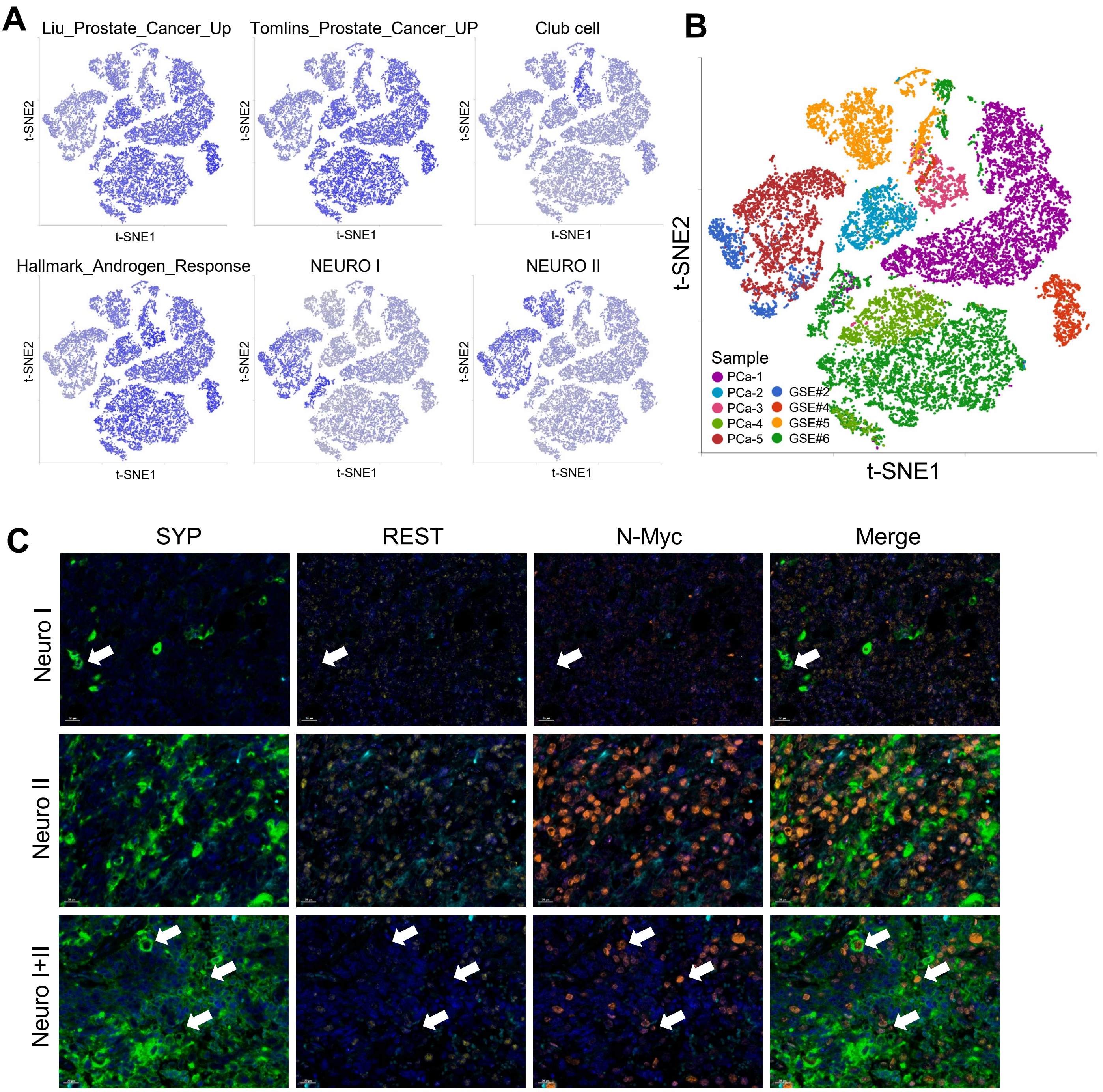

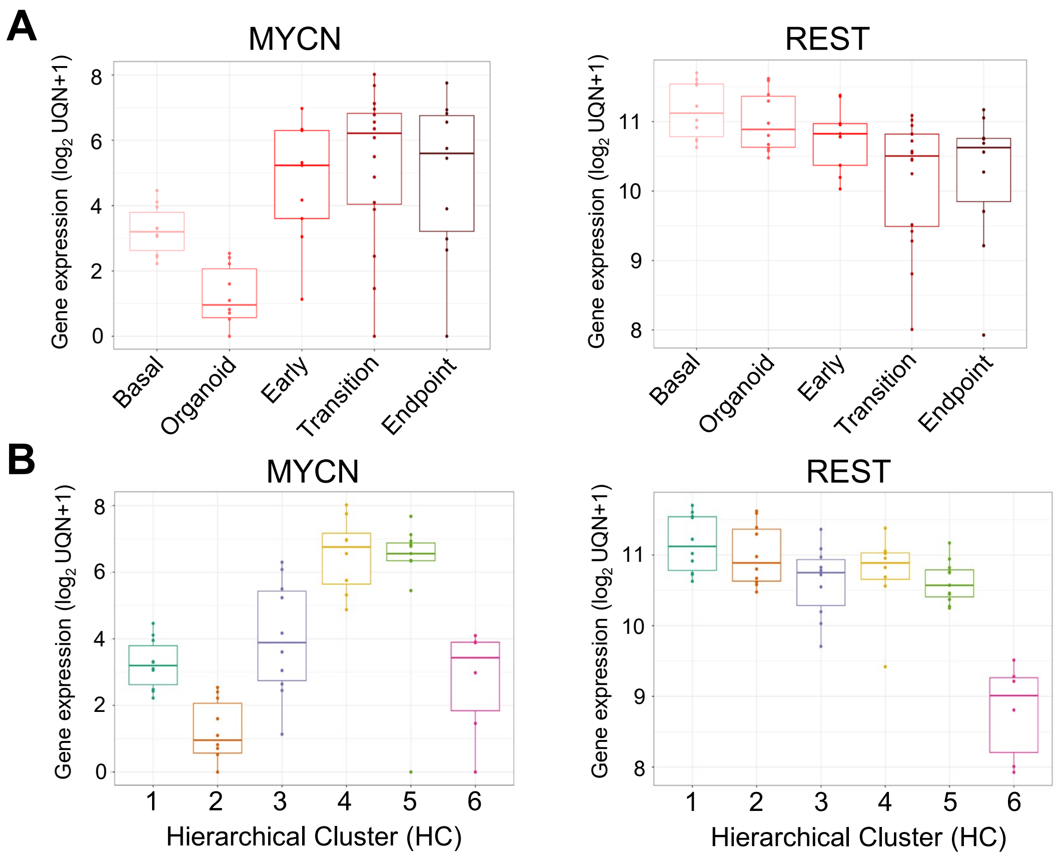

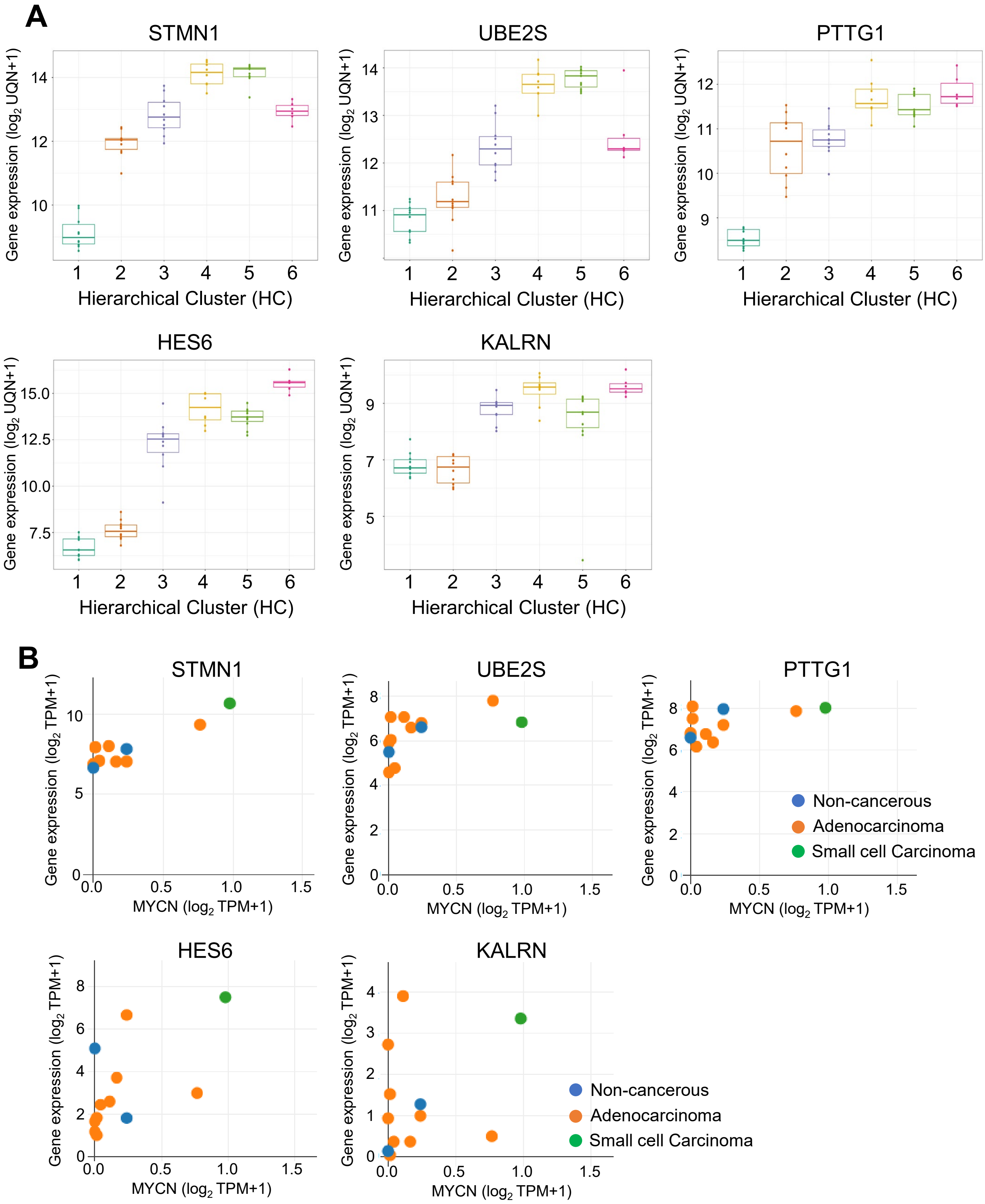
